# Pathogenicity, tissue tropism and potential vertical transmission of SARSr-CoV-2 in Malayan pangolins

**DOI:** 10.1101/2020.06.22.164442

**Authors:** Xiaobing Li, Kangpeng Xiao, Xiaoyuan Chen, Xianghui Liang, Xu Zhang, Zhipeng Zhang, Junqiong Zhai, Ruichen Wang, Niu Zhou, Zu-Jin Chen, Renwei Su, Fuqing Zhou, Edward C. Holmes, David M. Irwin, Rui-Ai Chen, Qian He, Ya-Jiang Wu, Chen Wang, Xue-Qing Du, Shi-Ming Peng, Wei-Jun Xie, Fen Shan, Wan-Ping Li, Jun-Wei Dai, Xuejuan Shen, Yaoyu Feng, Lihua Xiao, Wu Chen, Yongyi Shen

## Abstract

SARS-CoV-2 is having severe impact on public health at a global scale. Malayan pangolin SARS-CoV-2-related coronavirus (SARSr-CoV-2) is closely related to SARS-CoV-2. We show that CT scans of virus-positive pangolins reveal bilateral ground-glass opacities in lungs in similar manner to COVID-19 patients. The virus infected multiple organs in pangolins, with the lungs being the major target. Histological expression showed that ACE2 and TMPRSS2 are co-expressed with viral RNA. Transcriptome analysis revealed an inadequate interferon response, with different dysregulated chemokines and cytokines responses in pregnant and non-pregnant adults and fetuses. Viral RNA and protein were detected in three fetuses providing evidence for vertical virus transmission. In sum, our study identifies the biological framework of SARSr-CoV-2 in pangolins, revealing striking similarities to COVID-19 in humans.

---

The ongoing pandemic of COVID-19 caused the coronavirus SARS-CoV-2 is severely effecting global health (*1–3*). After SARS-CoV and MERS-CoV, SARS-CoV-2 is the third coronavirus to cause severe respiratory illness in humans identified in the past two decades, although with a markedly lower infection fatality ratio (*4*). SARS-CoV-2 belongs to the genus *Betacoronavirus*, sharing 79.5% overall genome nucleotide sequence identity with SARS-CoV and 96.2% with the bat-derived SARS-related coronavirus RaTG13 to which it is most closely related at the level of the whole genome (*5*). A better understanding of the pathobiology of SARS-CoV-2 is critical for its control and the development of therapeutics. Several animal models are being developed to assist in better understanding of the pathogenicity of SARS-CoV-2. These include the development of transduced or transgenic mouse models that express human ACE2 and can be infected by SARS-CoV-2 (*6–8*), and macaques that can be infected and recapitulate moderate disease (*9, 10*).

We previously isolated a SARS-CoV-2-related coronavirus (SARSr-CoV-2) in Malayan pangolins (*Manis javanica*) that is closely related to SARS-CoV-2, especially in the receptor-binding domain (RBD) of the spike protein (*11, 12*). Understanding the biology of SARSr-CoV-2 in pangolins will be important for understanding key aspects of coronavirus disease.

## Identifying SARSr-CoV-2 infection

Herein, we studied 28 Malayan pangolins that were confiscated in Guangdong Province between March-August 2019 and initially sent to Guangzhou wildlife rescue center. Notably, seven of these animals were pregnant. We used the viral S (spike) gene to identify infection with the pangolin-derived SARSr-CoV-2, and found that 15 animals, including six pregnant females, naturally infected by the virus. Notably, the S genes amplified from these individuals exhibited no changes in the consensus sequence. Four samples were chosen for deep sequencing, enabling the *de novo* assembly of four near complete coronavirus genomes (Figure S1). Analyses of these genomes revealed that they exhibited 99.6-99.9% genetic identity with the pangolin SARSr-CoV-2 reference genome (EPI_ISL_410721) and hence are indicative of a localized outbreak due to a single source of infection (Figure S2).

## Pathogenicity of SARSr-CoV-2 in Malayan pangolins

COVID-19 is characterized by a range of symptoms, but in most cases include fever, cough, dyspnoea, and myalgia (*13*). Bilateral opacities on x-ray or patchy shadows and ground glass opacities visible in CT scans are the most common features of severe COVID-19 cases (*1, 14*). Malayan pangolins that were positive for pangolin SARSr-CoV-2 exhibited respiratory symptoms such as cough and shortness of breath (*11*). Blood gas tests revealed elevated levels of PCO_2_ (58.2, and 60.25 mmHg), HCO_3_ (29.9, and 34.6 mmol/L), and TCO_2_ (31.5 mmol/L) in the infected animals (Table S1), indicative of dyspnea. CT scans in one virus positive pangolin showed a diffused bilateral distribution of ground-glass opacities in the lungs (Figure 1A), similar to early-phase COVID-19 pneumonia. The other virus-positive pangolin exhibited air bronchogram, fibrotic streaks and a subpleural transparent line (Figure 1B), similar to the advanced phase of COVID-19 disease (*15*). Hence, pangolins naturally infected with SARSr-CoV-2 show similar symptoms and CT features as COVID-19 pneumonia in humans.

**Figure 1.**
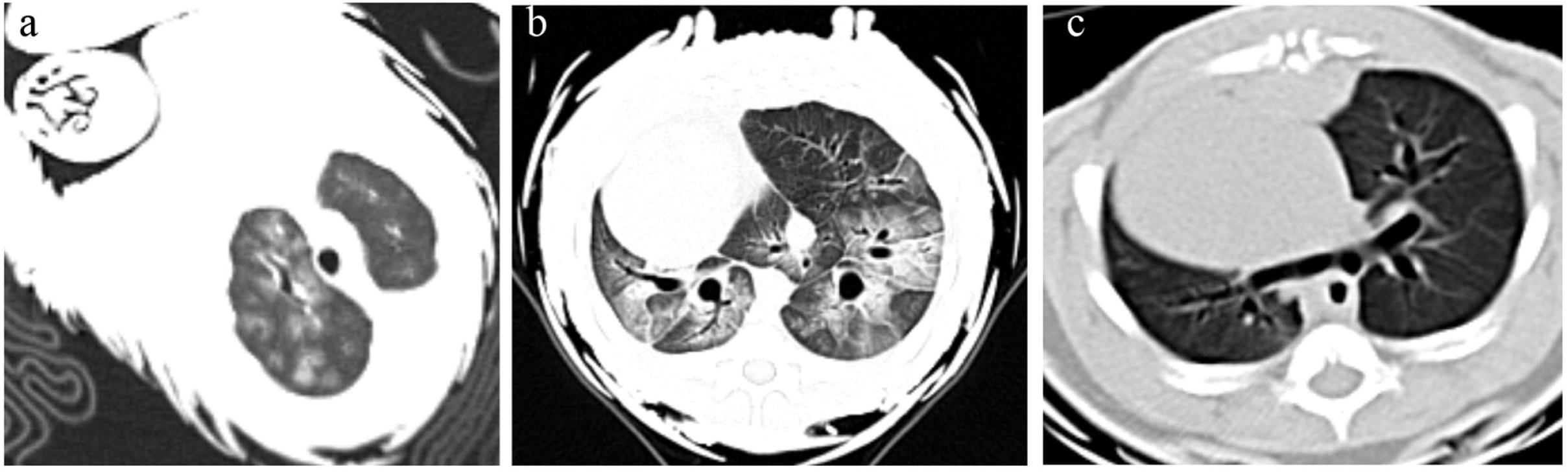
Chest CTs of two SARSr-CoV-2-positive and one negative pangolin. (a) Multiple ground-glass opacities in bilateral lungs in one virus-positive pangolin. (b) Bilateral focal consolidation, lobar consolidation, and patchy consolidation were clearly observed in the other virus-positive pangolin. (c) Chest CT of a virus-negative pangolin.

Necropsy showed pulmonary edema and congestion in lungs of virus-positive pangolins (Figure S3). Lungs from six pangolins, including four were positive for SARSr-CoV-2, were stained with hematoxylin and eosin (H&E) for histological examinations. Compared with virus-negative pangolins, lung tissues from the virus-positive animals showed severe interstitial pneumonia, in which the interstitium and the walls of the alveoli were thickened and displayed bronchiectasis (Figure 2A-C and Figure S4). These features are consistent with breathing difficulties. Alveoli were filled with desquamated epithelial cells and some macrophages with hemosiderin pigments (Figure 2C).

**Figure 2.**
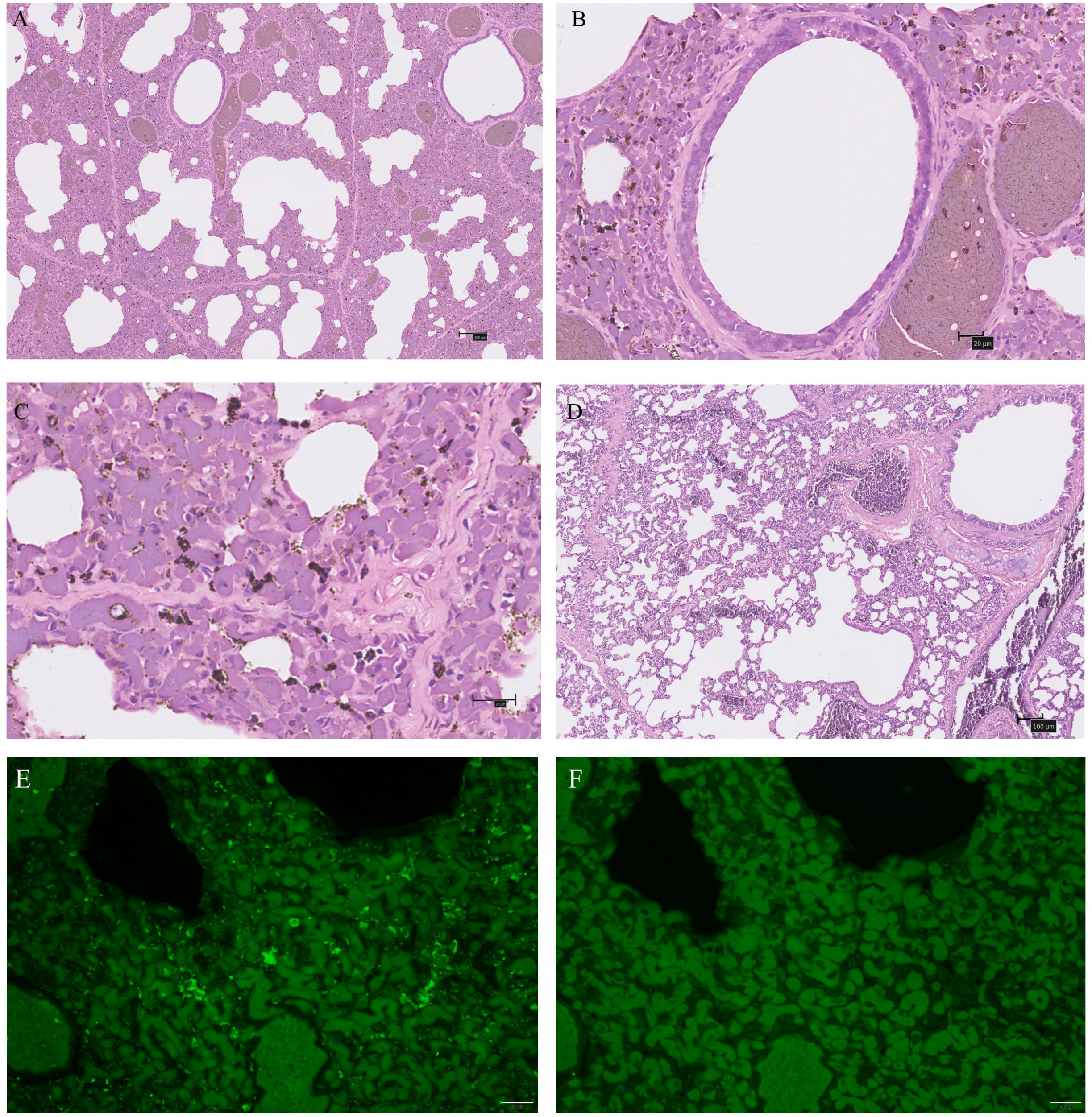
Characteristic pathological changes and virus antigen expression in the lungs of one SARSr-CoV-2 positive pangolin. (A-C) An autopsy lung with SARSr-CoV-2 infection compared to a negative one (D) were stained by hematoxylin and eosin (H&E). (B) bronchiectasis; (C) the interstitium and the walls of the alveoli thicken. (E-F) Immunofluorescence for virus antigen expression in pneumocytes. (E) nucleocapsid of SARS-CoV-2 (Green); (F) negative control. Scale bars of (A) and (D) are 100um, while others are 20um.

To ensure that these lesions were associated with SARSr-CoV-2, we used immunohistochemistry to detect and localize the nucleocapsid (N protein) of the virus. Several alveolar macrophages and pneumocytes were found to be positive (Figure 2E). qRT-PCR and western blot analysis further supported the presence of SARSr-CoV-2 infection in the lungs (Figure 2E, Figure 3A-B). These findings are similar to those of SARS-CoV-2 in humans and are compatible with a similar pathological mechanism, mainly by invading alveolar epithelial cells which results in respiratory symptoms (*13*).

## Tissue tropism of SARSr-CoV-2

Although respiratory symptoms dominate the clinical presentation of COVID-19, SARS-CoV-2 is also associated with multiple organ dysfunction syndrome (*13*). Necropsy of virus-positive pangolins similarly revealed multiple organ injury (Figure S1). We therefore used qRT-PCR to examine the tissue tropism of SARSr-CoV-2. Viral RNA was mainly detected in the lungs, although other organs such as liver, intestine, heart, kidney, spleen and muscle of some individuals also had detectable levels of virus RNA (Figure 3A, Table S2). Western blotting further supported the presence of viral infection in these tissues (Figure 3B).

**Figure 3.**
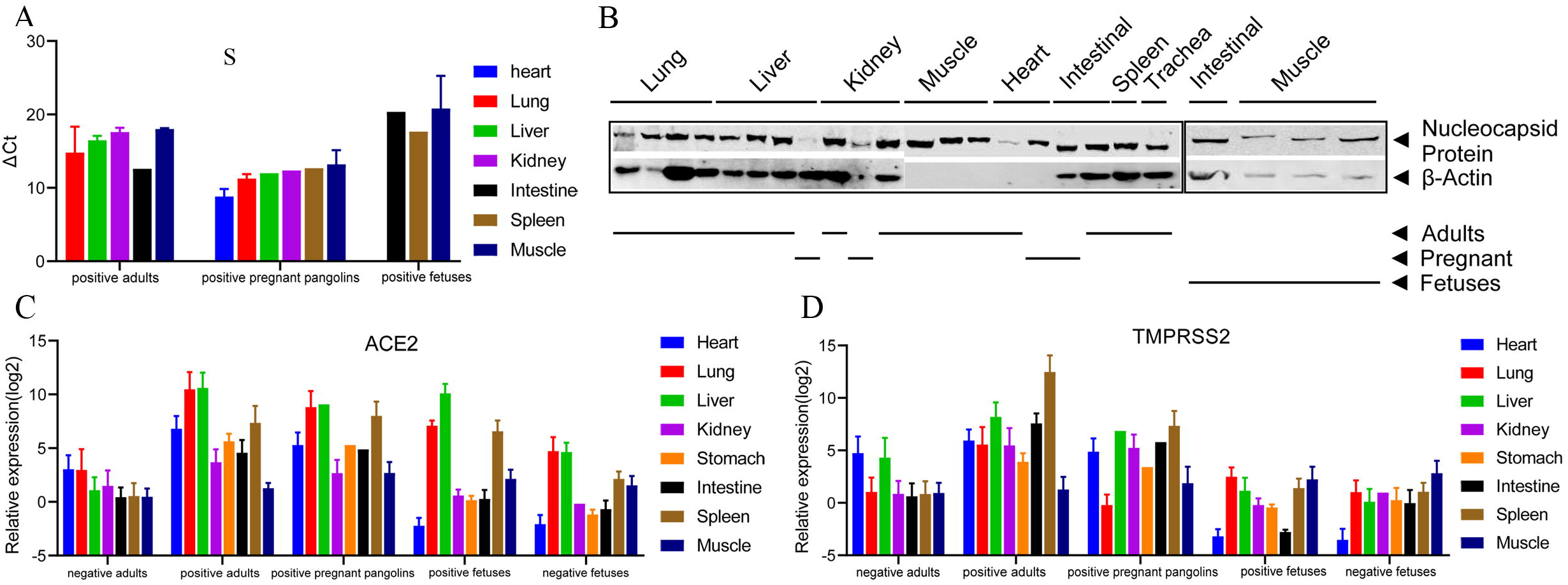
Expression patterns of the SARSr-CoV-2 and its receptor ACE2 and TMPRSS2 genes in different tissues of adult, pregnant pangolins and their fetuses. (A) Quantitative real-time PCR (qRT-PCR) of the S gene of SARSr-CoV-2. (B) Western blot of the N protein from the virus-positive tissues. Tissue lysates were subjected to SDS-PAGE and immunoblotting with antisera against SARS-CoV-2 N protein. (C) qRT-PCR of the ACE2 gene. (D) Quantitative real-time PCR (qRT-PCR) of the TMPRSS2 gene.

The spike protein of SARS-CoV-2 binds to ACE2 receptors for entry into cells (*5, 16*), while TMPRSS2 mediates spike protein activation and facilitates viral entry (*16, 17*). Generally, virus-positive pangolins had much higher expression levels of ACE2 and TMPRSS2 in all tissues than virus-negative individuals (Figure 3C and D). In humans, ACE2 is abundant in the epithelia of the lung, kidney, and small intestine (*18*). In pangolins, lung, liver, and spleen had the highest levels of ACE2 and TMPRSS expression compared to other tissues examined. SARSr-CoV-2 RNA was also detected in the tissues expressing the highest levels of ACE2 and TMPRSS, supporting the idea that cells that express both ACE2 and TMPRSS2 are most susceptible to coronaviruses (*19*). However, SARSr-CoV-2 was also detected in intestine and kidney, tissues that do not have high expression of these receptor genes in the pangolins.

## Potential vertical transmission of SARSr-CoV-2

As COVID-19 becomes a global pandemic, the potential risk of vertical transmission of SARS-CoV-2 has become a topic of concern (*20, 21*). The ACE2 receptor is widely expressed in the placenta during pregnancy (*22*). Hence, there is a theoretical risk of vertical transmission of SARS-CoV and SARS-CoV-2, both of which use ACE2 as receptor. Although neonatal cases of COVID-19 infection and antibodies in newborns have been found (*23–25*), testing was only performed in newborns (rather than embryos), so that infection after birth, or transfer of antibodies from the placenta and breast milk cannot be excluded. Whether vertical transmission of this virus occurs in humans therefore remains unclear and will ultimately require detection of the virus in embryos. We assessed the possibility of vertical virus transmission in seven pregnant pangolins. Of the six virus-positive pregnant pangolins, SARSr-CoV-2 RNA and N protein was detected in the intestine, spleen, and muscle of three of their fetuses (Figure 3A-B, Table S2). Hence, these results demonstrate it is possible that the vertical transmission of SARSr-CoV-2 can occur *in utero*.

## Host transcriptional response to SARSr-CoV-2

We next compared the transcriptional response of lung, spleen, muscle and intestine tissues to SARSr-CoV-2 infection between virus-positive versus virus-negative pangolins (Figure 4, Figures S5-6, and Table S3). Notably, up-regulated differentially expressed genes (DEGs) were enriched in lactate transport in lungs of virus-positive adults, likely reflecting hypoxia caused by pulmonary interstitial pneumonia. Similar to human tissues infected with SARS-CoV-2 or SARS-CoV (*26, 27*), a gene enrichment analyses illustrated that SARSr-CoV-2 triggered a dysregulated immune system in pangolins (Figure 4). Notably, non-pregnant adult, pregnant pangolins and fetuses had very different immune responses to the virus (Figure 4). In spleen, virus-positive fetuses showed evidence of the suppression of immune pathways: down-regulated DEGs were those associated with lymphocyte activation, leukocyte activation, interferon-gamma production, innate immune response, and cytokine production. For example, gene members of the TNF-receptor superfamily, such as CD3D, CD3E, CD3G, CD6, CD7, CD8A; CD8B, CD84, and CD247 were significantly down-regulated in spleen of fetuses. In contrast, pregnant pangolins showed up-regulated genes directed toward the regulation of humoral immune response, regulation of complement activation, inflammatory responses, and humoral immune responses in spleen, while adult pangolins had up-regulated genes enriched in negative regulation of type I interferon (IFN) production. The fetal immune system is unique and immature (*28*). Fetuses had no detectable virus RNA in lungs, and DEGs related to immune pathways were not enriched in lungs (Figure 4D). In the spleen and intestine, where virus RNA was detected in fetuses, down-regulated genes were related to immune pathways (Figure 4D and Figure S6).

**Figure 4.**
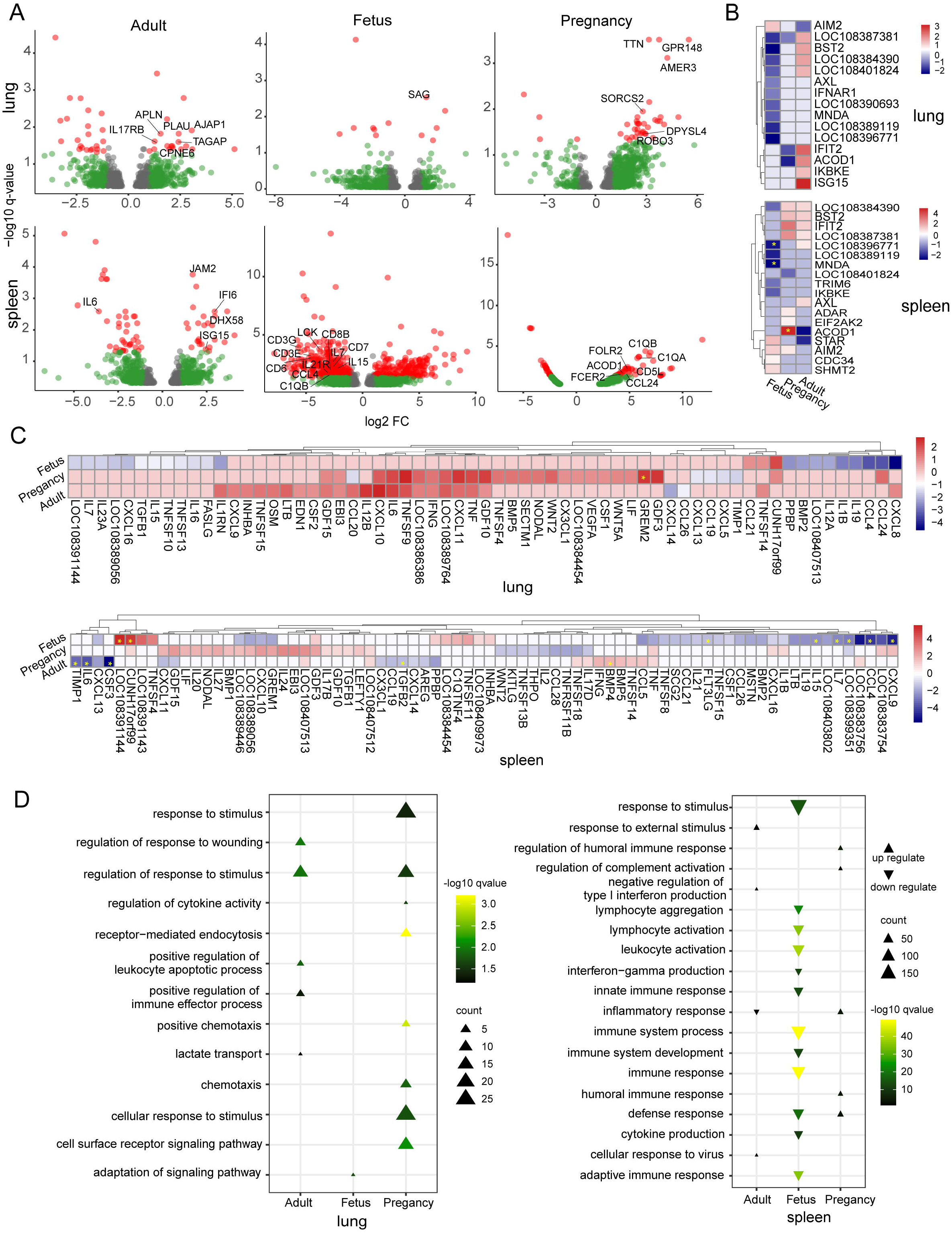
Host transcriptional response to SARSr-CoV-2 in the lung and spleen. (A) Volcano plot of DEGs comparing virus-positive versus virus-negative pangolins. (B) Heatmap of DEGs belonging to ISGs. (C) Heatmap of DEGs belonging to cytokine-related genes. (D) Functional enrichment analysis of DEGs.

After cellular detection of viral entry into a host cell, interferon induction of ISGs is essential for the host antiviral response (*29, 30*). Similar to SARS-CoV and SARS-CoV-2 (*26, 31*), we found only a small number of significant increases of ISGs expression in virus-positive pangolins in all tissues, implying an inadequate interferon response (Figure 4B, Figure S5B and S6B). For example, in lungs, IFIT2, ISG15, ACOD1, BST2 and IKBKE are up-regulated in adults, while there were down-regulated in fetuses and pregnant females. In spleen, BST2 and IFIT2 were up-regulated in adults and pregnant females, ACOD1 was down-regulated in adults and fetuses, but up-regulated in pregnant animals. ISG15 was up-regulated in muscle of adults and pregnant females but down-regulated in fetuses, while AIM2 showed the reverse trend (Figure S5). BST2 was significantly down-regulated in the intestine of virus-positive fetuses (Figure S6). Some coding proteins of SARSr-CoV-2, such as N and orf6, may therefore inhibit IFN1 by regulating IFN-β synthesis and signaling to avoid the innate immune response in a similar manner to other coronaviruses (*32–34*).

Many of the transcriptomic profiling studies of COVID-19 have been conducted using peripheral blood, often revealing the upregulation of chemokines and cytokines (*35*). Transcriptomic profiles of virus-positive pangolins differed among lung, spleen, muscle and intestine, while only the lung, the major target of infection, exhibited a greater signals of both cytokine and chemokine activity (Figure 4, Figure S5 and S6). Interleukin (IL) 1 Beta, IL6, IL7, IL12A, IL12B, IL15, IL16, IL19, IL23A, C-C Motif Chemokine Ligand (CCL) 4, CCL24, CCL21, C-X-C Motif Chemokine Ligand (CXCL) 8, CXCL9, CXCL10, CXCL11, CXCL16, TNF superfamily member (TNFSF) 4, TNFSF9, TNFSF10, TNFSF13, TNFSF14, TNFSF15 were up-regulated in lungs in both adults and pregnant females. Innate immunity can lead to an excessive inflammatory response and an over-exuberant cytokine response, with IL-6 having a direct association with the severe patient condition with SARS-CoV-2 and SARS-CoV (*36, 37*). Expression of CXCL8 (IL-8) is an attractant for neutrophils and associated with SARS-CoV-2 infection (*26*). Similarly, we found up-regulated expression of IL-6 and CXCL8 genes in both adults and pregnant females.

In sum, we show that Malayan pangolins infected with SARSr-CoV-2 exhibit a strikingly similar pathogenicity to human COVID-19 patients. For example, the host response of pangolins to SARSr-CoV-2, reflected in a low level ISG response, is similar to that seen in human COVID-19. Although the SARS-CoV-2-related pangolin virus has a number of important genetic differences to SARS-CoV-2, we were able to show that SARSr-CoV-2 was present in pangolin fetuses highlights the potential for *in utero* virus transmission.

## ACKNOWLEDGMENTS

We thank Dr. Zhengli Shi, Wuhan Institute of Virology, for providing SARS-CoV-2 nucleocapsid antibody.

## Author contributions

Y.S. conceived the project; W.C., J.Z., N.Z., Z-J. C., Y-J. W., C. W., X-Q. D., S-M. P., W-J. X., F. S., W-P. L., and J-W. D. performed the pangolin necropsy and tissue collection; W.C., and F.Z. performed the CT scans and blood tests. R.S. undertook the pathological examinations. X.C., Q.H., and X.L. performed the qRT-PCR; X.Z., X.C., and X.S. carried out the immunofluorescence experiments; X.L., Z.Z., and R.W. performed western blots; K.X., X.L., E.C.H., and Y.S. contributed to the transcriptome analyses and interpretation; the manuscript was written by Y.S., E.C.H., D.M.I., and R.A.C. All authors discussed the experiments and results, read and approved the manuscript.

## Data and materials availability

The raw sequencing datasets generated during this study are available on the NCBI Gene Expression Omnibus (GEO) server under the accession number PRJNA640246. The S gene sequences and near complete genome sequence of pangolin-derived SARSr-CoV-2 has been deposited in GISAID with the accession numbers EPI_ISL_471461-EPI_ISL_471470.

## Supplementary Materials

### Materials and Methods

#### Samples

The Malayan pangolin (*Manis javanica*) is classified as critically endangered in accordance with the International Union for Conservation of Nature (IUCN) Red List of Threatened Species R, and included in Appendix I of the Convention on International Trade in Endangered Species of Wild Fauna and Flora (CITES I). The 28 adult Malayan pangolins, seven of which were pregnant, used in this study were confiscated by Customs and the Department of Forestry of Guangdong Province in March-August 2019, and naturally infected by the virus. The fetuses of the seven pregnant pangolins were also used. These animals were initially sent to Guangzhou wildlife rescue center, during with time some pangolins presented with respiratory symptoms. Computed tomographic (CT) scans were performed on two animals. For comparison, CT scan of another pangolin without respiratory symptoms was also performed. Unfortunately, despite exhaustive rescue efforts all of the pangolins eventually died. Tissues, including lung, liver, spleen, muscle, kidney, intestine, and heart were collected, and stored in −80°C refrigerator. As these pangolins died in 2019, prior to the emergence of COVID-19, the importance of these samples was not fully realized at the time so that sample collection was incomplete. Available tissue samples are shown in Table S2.

#### qRT-PCR analysis

SARSr-CoV-2 viral RNA, as well as ACE2 and TMPRSS2 transcript levels, were measured by real-time PCR using SYBR Green chemistry. Total RNA was isolated using the QIAamp Viral RNA Mini Kit (Qiagen) according to the kit instructions. 5μg of Total RNA was reverse transcribed into cDNA in a 20 μL reaction volume using by PrimeScript ™ IV 1st strand cDNA Synthesis Mix (Takara). For quantitative real-time PCR (qRT-PCR), cDNA was used with ChamQ™ Universal SYBR® qPCR Master Mix (vazyme) in a CFX Connect™ Real-Time System (BIO-RAD). Glyceraldehyde-3-phosphate dehydrogenase (GAPDH) was selected as a host reference gene to normalize the relative viral loads, evaluated by calculating -ΔΔCt values. Each quantitative PCR was performed in triplicate. The primers are shown in Table S4.

#### Western blots

Protein was extracted with radio-immuno precipitation assay (RIPA) buffer (50mM Tris, pH 8.0, 150mM NaCl, 1.0% NP-40, 0.5% sodium deoxycholate, 0.1% SDS) and a protease inhibitor cocktail. 10mg of total protein was separated on SDS-PAGE and then transferred to PVDF membranes using the Bio-Rad Trans-Blot protein transfer system. The membrane was blocked at room temperature for 90 min with 5% skim milk in 1 × TBST (20 mM Tris–HCl, pH 7.4, 150 mM NaCl, and 0.1% Tween 20). After blocking, the membrane was incubated with SARS-CoV-2 nucleocapsid antibody (kindly provided by Dr. Zhengli Shi, Wuhan Institute of Virology), and Recombinant Anti-Actin antibody (Abcam) at 4°C overnight, washed three times with 1 × TBST, and incubated with corresponding peroxidase-conjugated (Thermo Fisher Scientific) or fluorescently labeled (IRDye 680RD or 800CW: Li-Cor Biosciences) secondary antibody. The membranes were washed three times with 1 × TBST, and then scanned on an Odyssey Infrared Imaging System (Li-Cor Biosciences).

#### Histopathology and immunofluorescence

Lungs from six Malayan pangolins, including four that were positive for SARSr-CoV-2, were fixed in 10% buffered formalin. Tissues were then washed to remove formalin, dehydrated in ascending grades of ethanol, cleared with chloroform, and embedded with molten paraffin wax in a template. The tissue blocks were sectioned with a microtome, and sections were transferred onto grease-free glass slides, deparaffinized, and rehydrated through descending grades of ethanol and distilled water.

Sections were stained with a hematoxylin and eosin staining kit (Baso Diagnostics Inc., Wuhan Servicebio Technology Co., Ltd.). Other sections were incubated in 3% H2O2 solution in methanol to block endogenous peroxidase activity after dewaxing in xylene and hydrating in different concentrations of ethanol (100%, 95%, 85%, 75% and 50%). Subsequently, sections were incubated in permeabilizing solution containing 1% Triton X-100 (Beijing Solarbio Science & Technology Co., Ltd) in PBS for one hour and then treated with microwaves for antigen retrieval and washed with PBS. Sections were then blocked using 5% BSA at room temperature for one hour and incubated overnight in a humidified chamber with primary antibody against SARS-CoV-2 nucleocapsid (1:100, kindly provided by Dr. Zhengli Shi) at 4°C. Following washing three times in PBS, sections were incubated with Biotin-goat anti-rabbit secondary antibody (1:100, Concentrated DyLight 488-SABC kit, Boster Biological Technology Co. Ltd, China) in a humidified chamber for 30 min at 37°C and then incubated with DyLight 488-SABC (1:200, Concentrated DyLight 488-SABC kit, Boster Biological Technology Co. Ltd, China) for 30 min at 37°C. After washing four times with PBS, sections were mounted with Antifade Mounting Medium with DAPI (Beijing ZSGB Biotechnology Co., Ltd). Tissue fluorescence was visualized using a fluorescence microscope (OLYMPUS, Tokyo, Japan).

#### RNA sequencing

TRIzol (Invitrogen) was used for the extraction of total RNA, with DNA digested with DNase I before library construction. tRNA was also removed before library preparation. Sequencing libraries were generated using NEBNext® Ultra™ RNA Library Prep Kit for Illumina® (NEB, USA) following the manufacturer’s recommendations and index codes were added to attribute the sequences to each sample. Sequencing libraries were sequenced on an Illumina NovaSeq 6000 platform, generating 150 bp paired-end reads. Approximately 12 Gb of raw data were generated for each sample (Table S3). The raw sequencing data sets were submitted to NCBI Gene Expression Omnibus (GEO) server (accession number PRJNA640246).

#### Bioinformatic analyses

Adaptor and low-quality sequences were trimmed using fastp (v0.19.7) (*38*). Viral sequences were counted by mapping the clean reads to the pangolin SARSr-CoV-2 genome (GIASID: EPI_ISL_410721) through BWA-MEM (v0.7.17) (*39*). Transcriptomes were *de novo* assembled using Megahit (v1.0.3) (*40*). The Malayan pangolin genome (ManJav1.0) was used as the reference host genome. Sequence alignments were undertaken using the program Hisat2 (*41*). Gene counts were summarized using the featureCounts program (*42*) as part of the Subread package release 2.0.0 (http://subread.sourceforge.net/). Differential expression analysis of two groups was performed using the DESeq2 R package (1.10.1) (*43*). The resulting p-values were adjusted using the Benjamini and Hochberg’s approach for controlling the false discovery rate. Differentially expressed genes (DEGs) were characterized for each sample (ļlog2 foldchangeļ > 1, p-adjusted-value < 0.05).

The Gene Ontology (GO) functional annotation of protein sequences were made on the basis of a BLASTp (v2.7.1+) alignment against the Swissprot protein database. The GO enrichment analysis of different expressed genes (DEGs) was performed using the clusterProfiler R package (*44*). Heatmaps of gene expression levels were constructing using pheatmap in R packages. Volcano plots and dolt plots were constructed using ggplot2 (*45*). The heatmap of Type-I IFN responses was constructed on genes with ļlog2 foldchangeļ > 1 belonging to the following GO annotations: GO:0035457, GO:0035458, GO:0035455, GO:0035456, GO:0034340. The heatmap of cytokine activity and chemokine activity was constructed on genes with |log2 foldchange| > 1 belonging GO annotations for GO: 0005125 and GO: 0008009.

---

**Figure S1.** Mapping of raw reads from the high throughput sequencing of four pangolin lung tissues to SARSr-CoV-2 reference genome (EPI_ISL_410721).

**Figure S2.** Phylogeny of coronaviruses closely related to SARS-CoV-2 based on full genome sequences. The phylogenetic tree was constructed using RAxML using the GTRGAMMAI substitution model and 1,000 bootstrap replicates. Numbers (>80) above or below branches are percentage bootstrap values for the associated nodes. The scale bar represents number of substitutions per site. Red circles indicate the pangolin coronavirus sequences generated in this study.

**Figure S3.** Photos taken soon before its death and at necropsy from a SARSr-CoV-2 positive Malayan. (a) Before its death; (b) Trachea hemorrhage; (c) Pulmonary edema, congestion, hemorrhage; (d) Nephromegaly and hemorrhage; (e) heart; (f) Hepatomegaly with obvious bleeding spots.

**Figure S4.** Histopathological changes in the lungs of three SARSr-CoV-2 positive pangolins. Scale bars are 100um.

**Figure S5.** Host transcriptional response of muscle to pangolin-CoV. (A) Volcano plot of DEGs comparing virus-positive versus virus-negative pangolins. (B) Heatmap of DEGs belonging to ISGs. (C) Heatmap of DEGs belonging to cytokine-related genes. (D) Functional enrichment analysis of DEGs.

**Figure S6.** Host transcriptional response of intestine of fetuses to pangolin-CoV. (A) Functional enrichment analysis of DEGs. (B) Volcano plot of DEGs comparing virus-positive versus virus-negative pangolins. (C) Heatmap of DEGs belonging to ISGs. (D) Heatmap of DEGs belonging to cytokine-related genes. Only intestines of fetuses are available.

**Table S1**. Blood gas analysis and routine blood tests of pangolins.

**Table S2**. ΔΔCt of qRT-PCR of S gene of SARSr-CoV-2, ACE2 and TMPRSS2 genes.

**Table S3**. Summary of pangolin transcriptomes.

**Table S4**. Primers used for qRT-PCR.

