## Supplementary figures and images for "Pathogenicity, tissue tropism and potential vertical transmission of SARSr-CoV-2 in Malayan pangolins"

### Figure S1

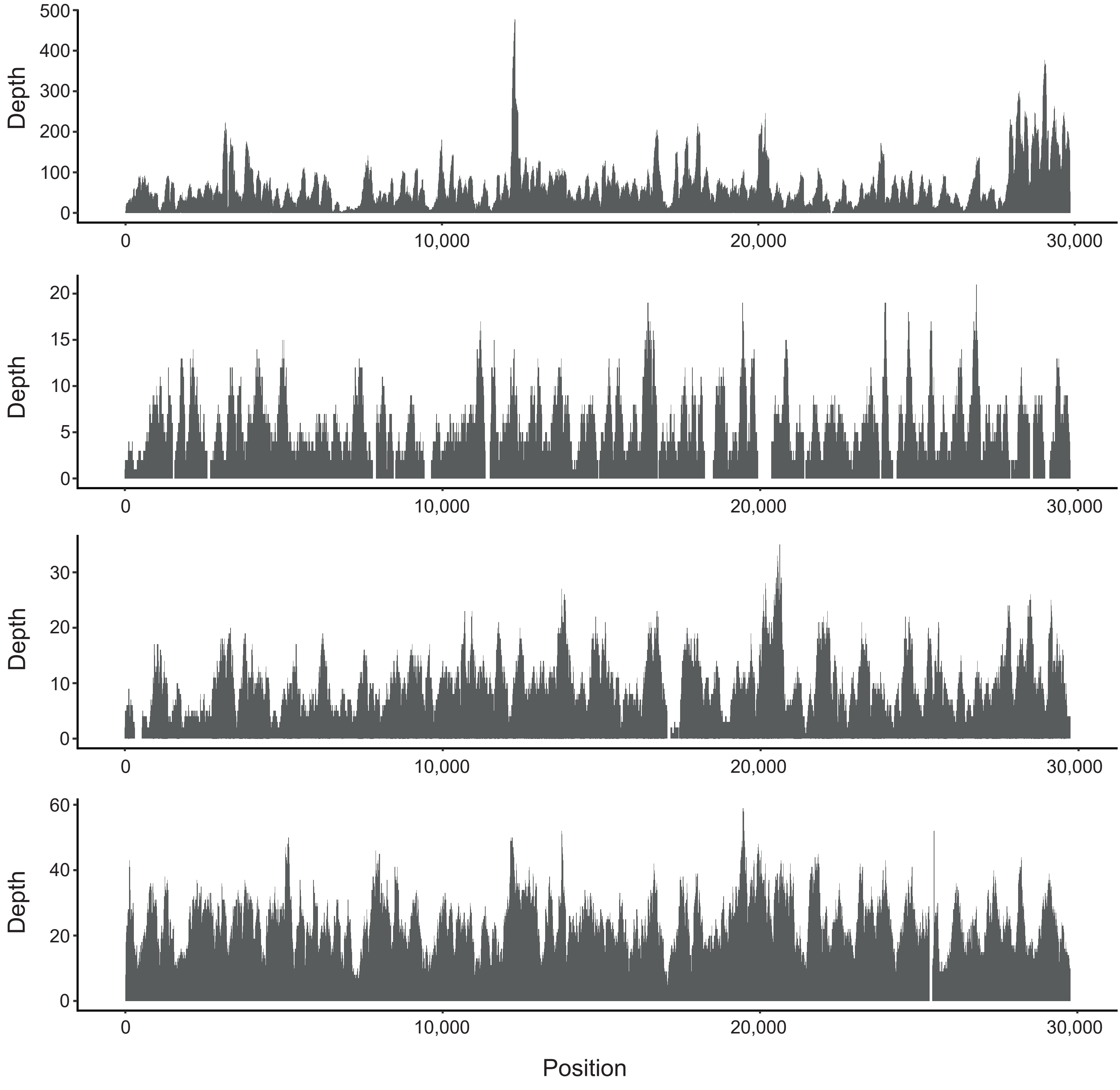

### Figure S2

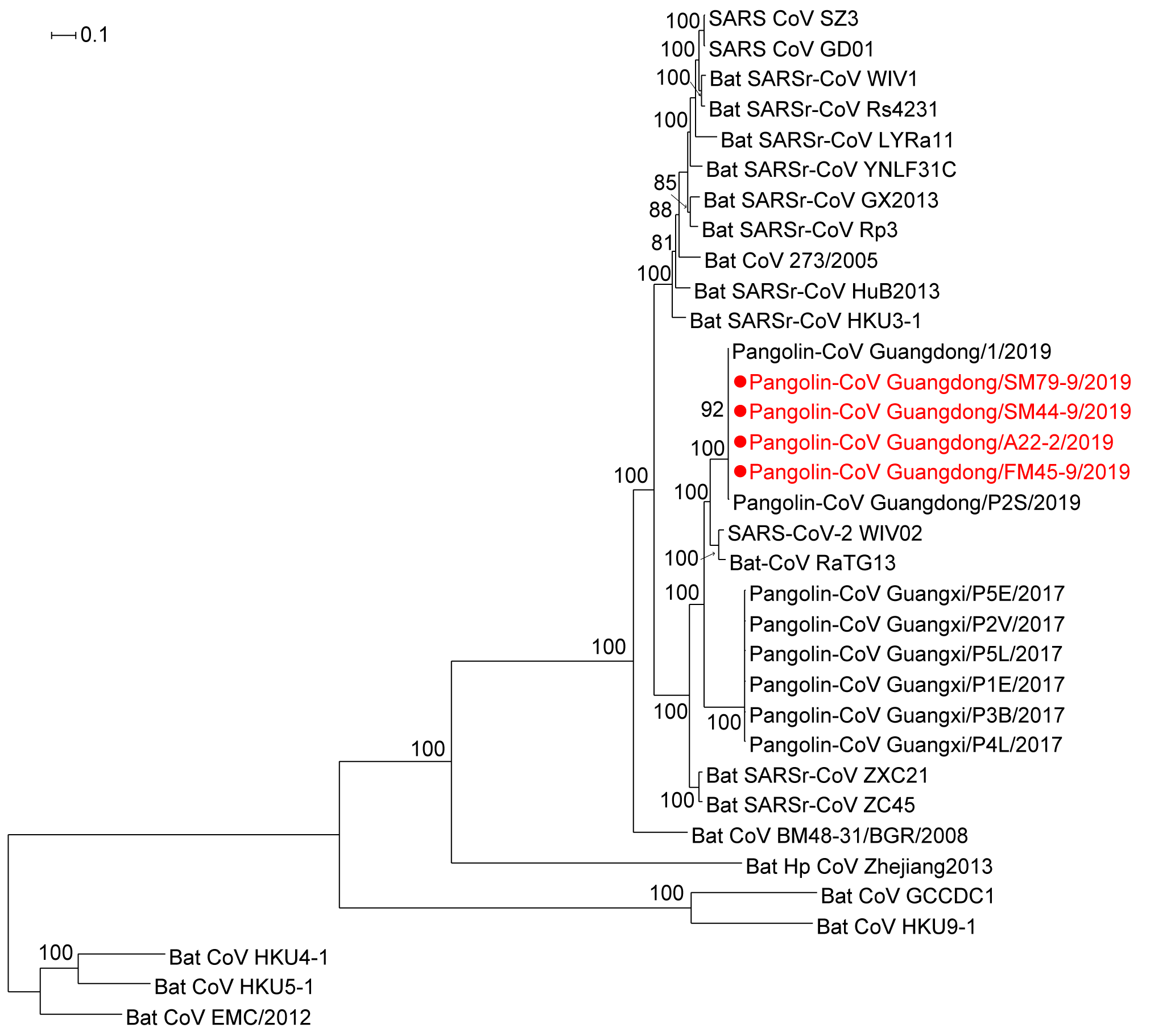

### Figure S3

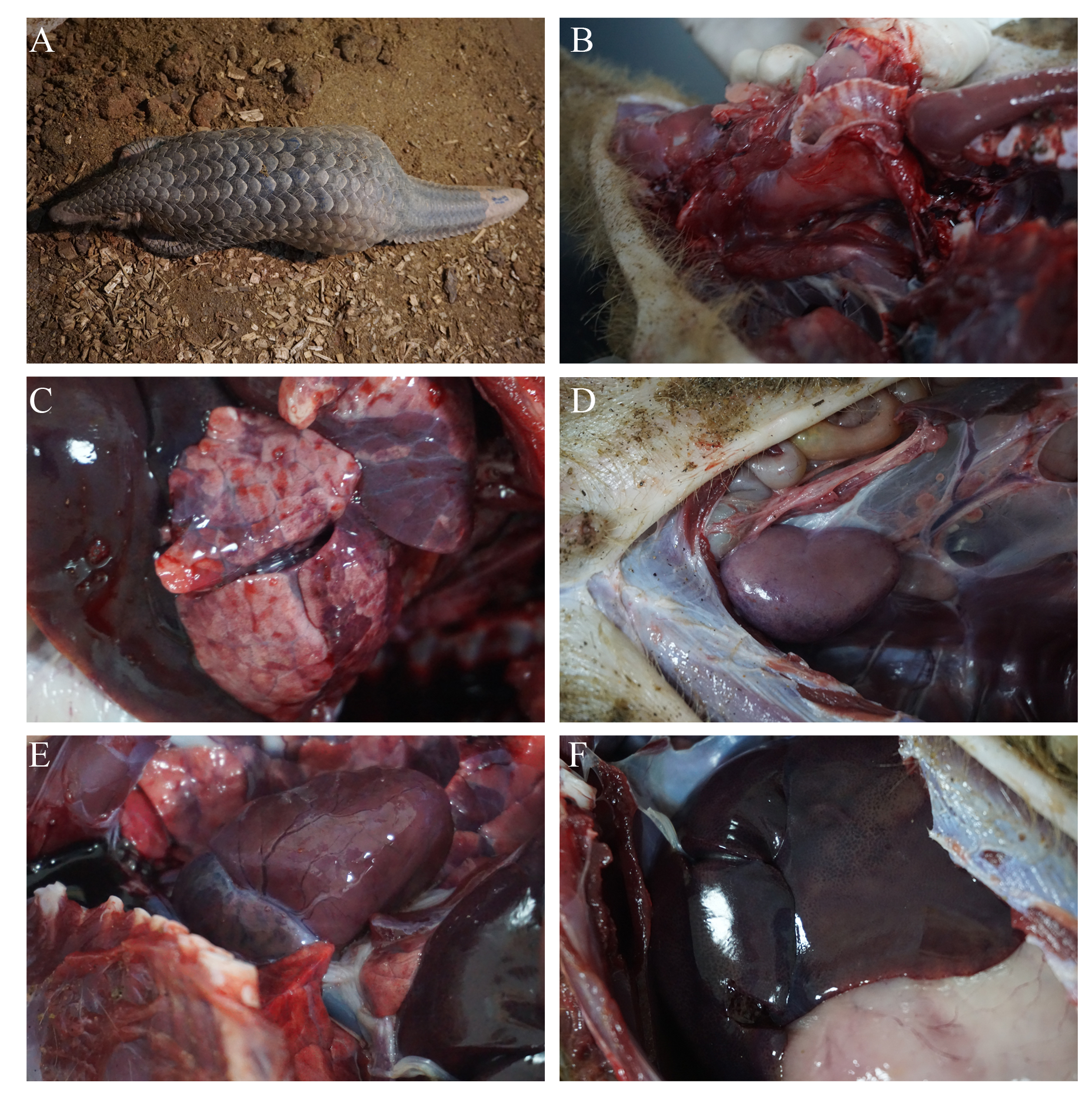

### Figure S4

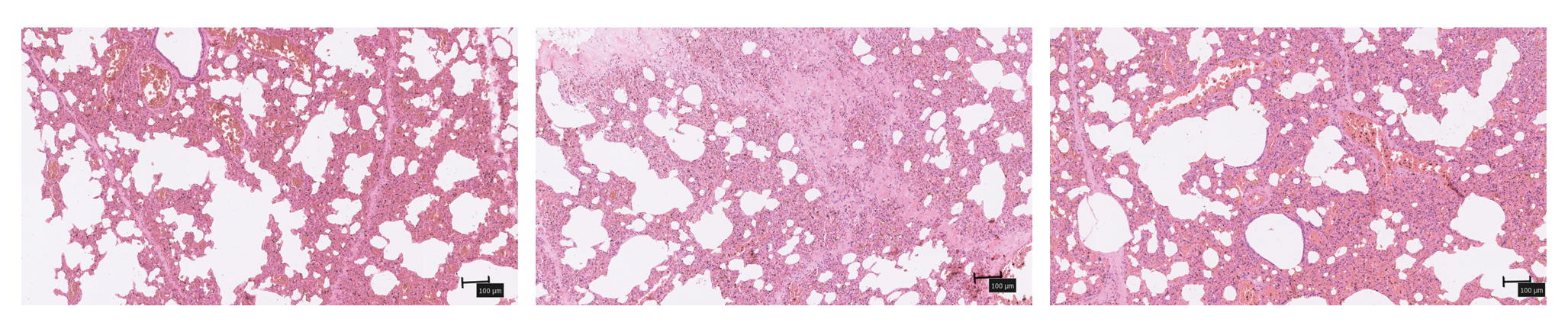

### Figure S5

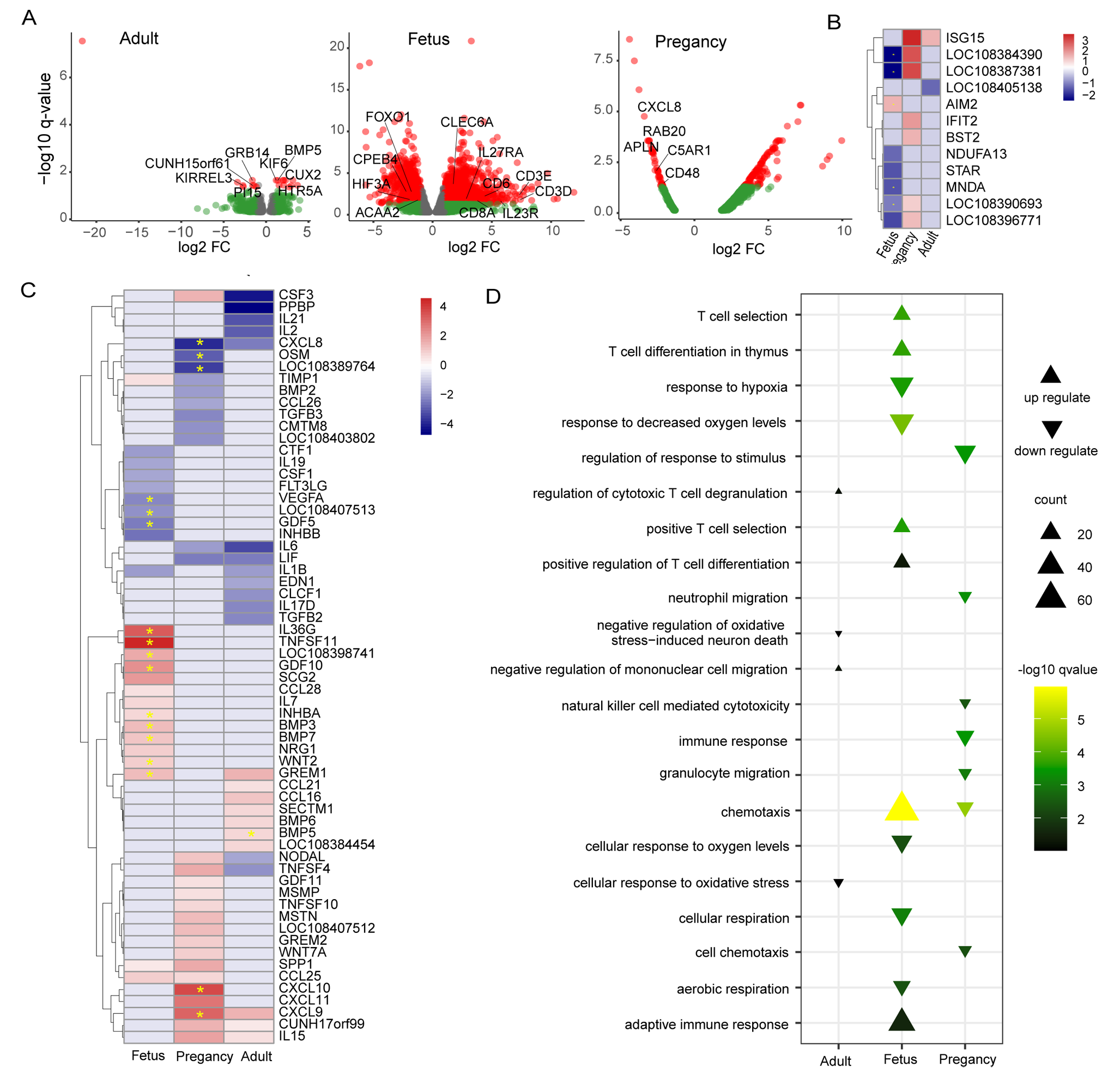

### Figure S6

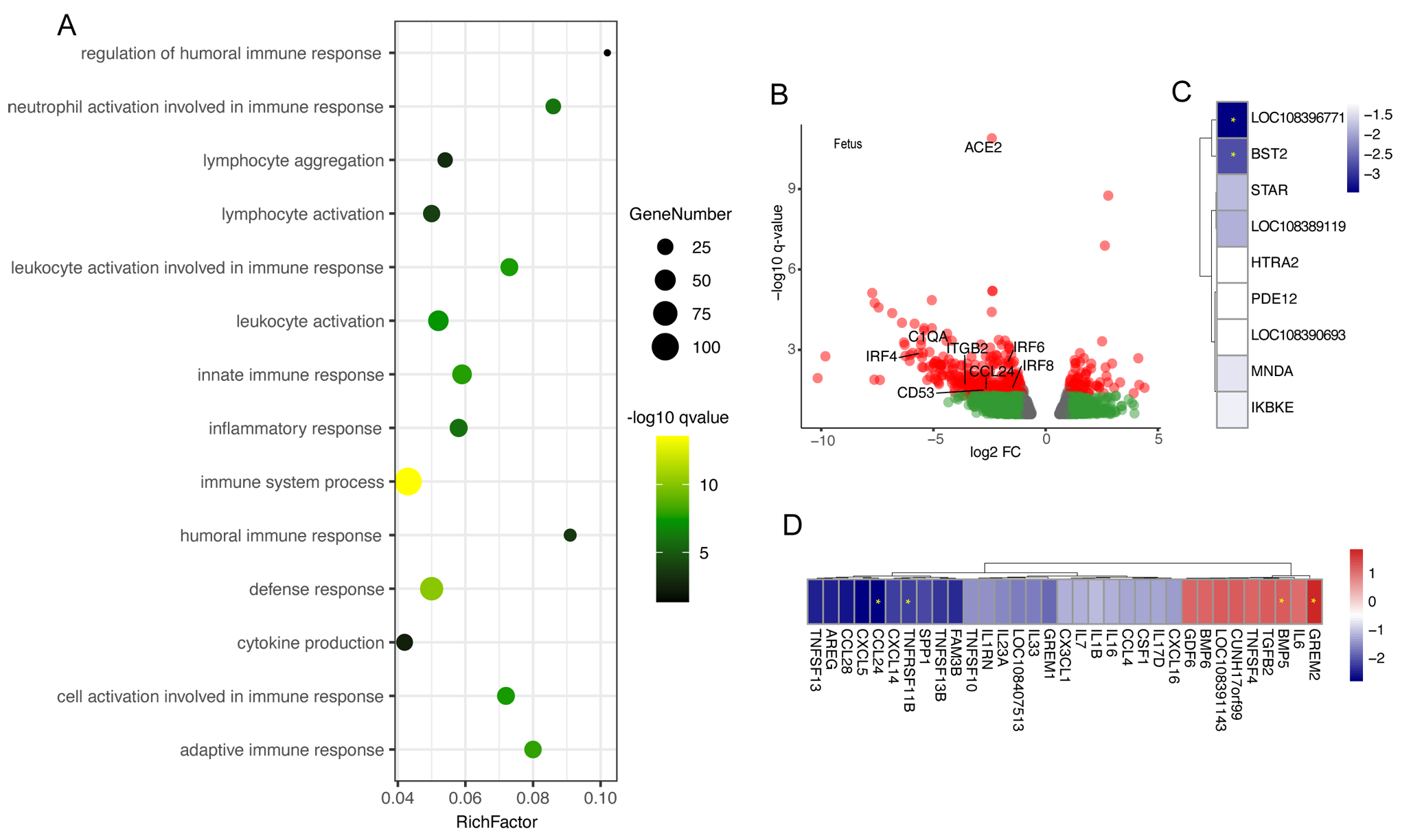
